# Learning of biased representations in LIP through interactions between recurrent connectivity and Hebbian plasticity

**DOI:** 10.1101/2021.09.23.461557

**Authors:** Wujie Zhang, Jacqueline Gottlieb, Kenneth D. Miller

## Abstract

When monkeys learn to group visual stimuli into arbitrary categories, lateral intraparietal area (LIP) neurons become category-selective. Surprisingly, the representations of learned categories are overwhelmingly biased: nearly all LIP neurons in a given animal prefer the same category over other behaviorally equivalent categories. We propose a model where such biased representations develop through the interplay between Hebbian plasticity and the recurrent connectivity of LIP. In this model, two separable processes of positive feedback unfold in parallel: in one, category selectivity emerges from competition between prefrontal inputs; in the other, bias develops due to lateral interactions among LIP neurons. This model reproduces the levels of category selectivity and bias observed under a variety of conditions, as well as the redevelopment of bias after monkeys learn redefined categories. It predicts that LIP receptive fields would spatially cluster by preferred category, which we experimentally confirm. In summary, our model reveals a mechanism by which LIP learns abstract representations and assigns meaning to sensory inputs.

## Introduction

The lateral intraparietal area (LIP) is an area of the monkey brain that interfaces between visual perception, eye movement planning, and visuospatial cognition. Neurons in LIP have visuospatial receptive fields (RFs), and it is well-known that they can encode visual salience, attention, saccadic eye movement, and visuospatial decision making variables (Bisley and Goldberg, 2010; Andersen and Cui, 2009; Gottlieb et al., 2009; Gold and Shadlen, 2007; Kable and Glimcher, 2009; Gottlieb, 2007). Moreover, LIP has been shown to encode non-spatial variables (Gottlieb and Snyder, 2010). For example, Oristaglio et al. (2006) showed that LIP neurons encoded whether the monkey used its left or right hand in an operant response. The non-spatial variables encoded by LIP can also be abstract: Freedman and Assad (2006) and Fitzgerald et al. (2011) found that, when monkeys learned to group visual stimuli into abstract categories, LIP neurons encoded these categories.

Surprisingly, this neural representation of categories carried a strong bias. Typically, in a given brain area, among neurons responsive to a set of stimuli, their preferred stimuli tend to evenly span the stimulus space (e.g., Hubel and Wiesel, 1962; DeAngelis and Uka, 2003; Conway and Tsao, 2009; Hegde and Van Essen, 2007; Lehky et al., 2011). In LIP, however, the representations of learned categories were overwhelmingly biased: while different categories were behaviorally equivalent, nearly all LIP neurons in a given animal preferred the same category (Fitzgerald et al., 2013). Similar biases were also seen in a perceptual decision making task (Bennur and Gold, 2011; Fitzgerald et al., 2013).

Here we built a model of Hebbian plasticity acting on the synapses from PFC to LIP. We show that recurrent connectivity in LIP allows simple Hebbian plasticity to give rise to category selectivity and representational bias. In the following text, we give an overview of the experimentally observed category selectivity and bias, lay out considerations that lead to the premises of our model, and reproduce experimental findings through simulations. We then analyze the model to determine the underlying mechanisms, which leads to a number of experimental predictions, one of which—that LIP RFs should spatially cluster by preferred category—we also test and confirm. Together, this work reveals a potential general mechanism by which LIP learns abstract representations and assigns meaning to sensory inputs.

## Results

### Category selectivity and representational bias in LIP

Freedman and Assad (2006) and Fitzgerald et al. (2011) trained monkeys on match-to-category tasks (Fig. 1A), in which a number of different visual stimuli are grouped into arbitrary categories. On a given trial, a sample stimulus and a test stimulus were presented, separated by a time delay; the monkey’s task was to respond according to whether the sample and test stimuli belonged to the same category. In these tasks, LIP neurons exhibited a visual response during the presentation of the sample stimulus and a sustained response during the delay period after sample stimulus offset (Fig. 1B-D, illustrating three example neurons). During both the sample visual response and the delay response, neurons fired at different rates based on category of the sample stimulus (the two categories are represented by the colors red and blue throughout Fig. 1). Notably, single LIP neurons robustly encoded the category to which stimuli belonged, but not the identity of stimuli (Freedman and Assad, 2006).

**Figure 1.**
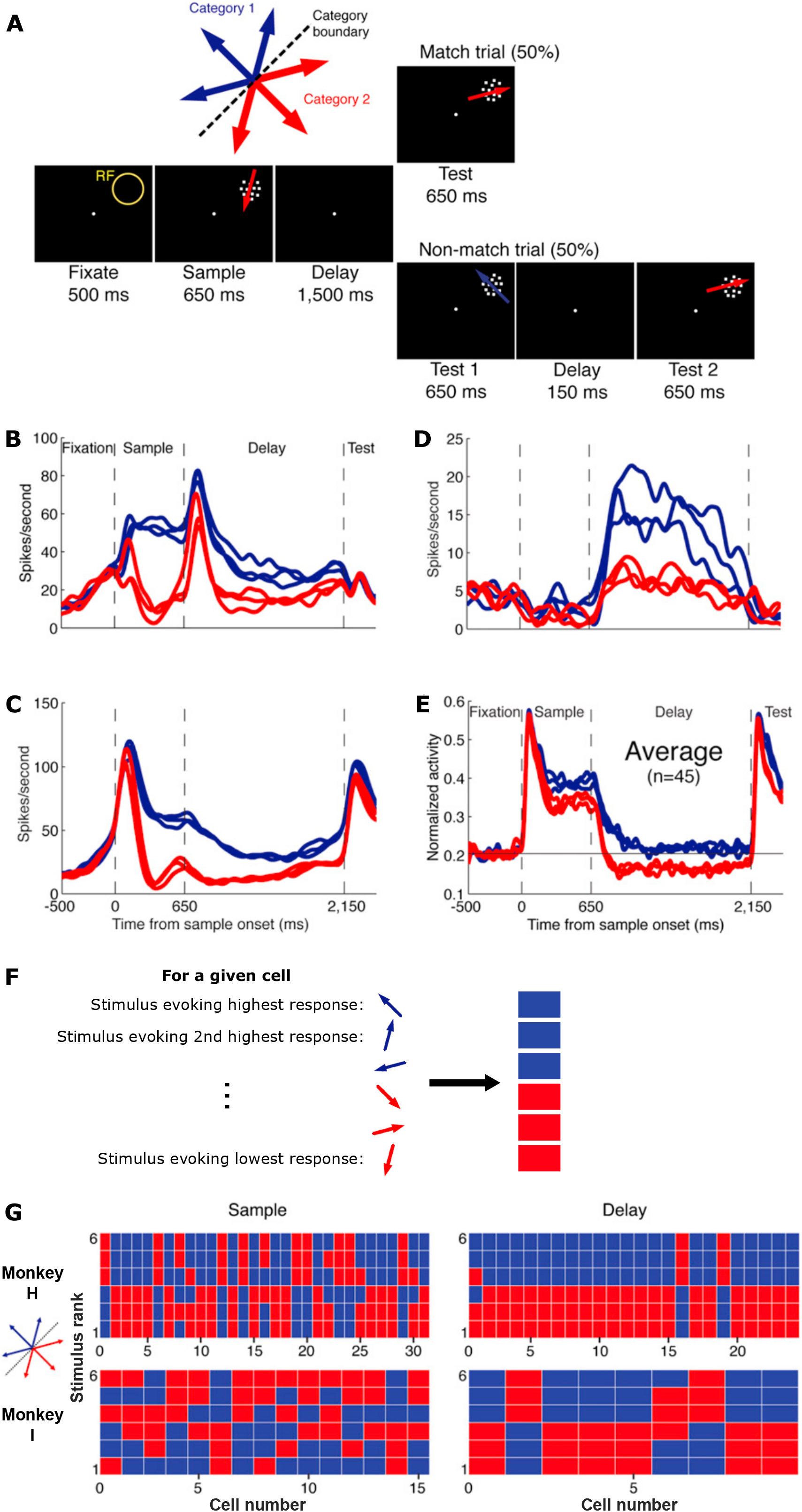
LIP shows biased representation of abstract categories. (A) The match-to-category task. After fixation, an array of coherently moving dots was shown as the sample stimulus inside the receptive field (RF). On a given trial, the sample stimulus could be moving in one of six directions, which were grouped into two categories. The sample stimulus was followed by a delay, after which another array of coherently moving dots was shown as the test stimulus. If the sample and the test stimuli belonged to the same category, the monkey was required to release a bar that it had been manually holding to receive a juice reward. If they belonged to different categories, the monkey was required to continue holding the bar until another test stimulus appeared that belonged to the same category as the sample. (B)-(D) PSTHs of three example cells from monkey H. In each panel, each of the six traces denotes trials in which one of the six sample stimuli was shown, colored according to the category to which the stimulus belonged. (E) Population average of normalized PSTHs from monkey H. Note that a category preference in the population average implies a biased representation among the cells. (F) Schematic illustrating a graphical representation of the rank ordering of responses for a single cell. The six sample stimuli are represented as rectangles that are colored according to their categories. The rectangles are arranged in a column, with the stimulus that evoked the highest response for the cell at the top, and the stimulus that evoked the second highest response second from the top, etc. (G) Rank ordering of neural responses during the sustained sample period (200-650 ms after motion onset; left panels) and late delay period (750-1500 ms after motion offset; right panels) for monkey H (top panels) and monkey I (bottom panels). In each panel, each column denotes a cell, and the ordering of stimuli is as explained in (F). Note that during the delay period, almost all cells preferred the blue category over the behaviorally equivalent red category. Figure adapted from Fitzgerald et al. (2013).

Examining the LIP population as a whole (Fig. 1E), we see that the average response of the population preferred one category over the other (firing more for the blue than the red category), suggesting that the different categories have unequal representation among the individual neurons. Indeed, an examination on a cell-by-cell basis shows that, especially during the delay period, most LIP neurons in a given monkey tended to have the same category preference (Fig. 1F-G; Fitzgerald et al., 2013). Furthermore, Fitzgerald et al. (2013) also observed a bias in the data of Bennur and Gold (2011). Bennur and Gold trained monkeys to report the direction of noisy arrays of moving dots, where leftward or rightward motion was reported by saccades to a red or green target, respectively. In two monkeys, LIP neurons overwhelmingly preferred rightward motion and red target over leftward motion and green target, during both the presentation of the motion stimuli and the delay period after their offset.

Thus, bias is a robust signature of categorical representation in LIP, being observed in three different experiments, in five monkeys from two laboratories (Fitzgerald et al., 2013; Freedman and Assad, 2006; Fitzgerald et al., 2011; Bennur and Gold, 2011). Such a biased coding scheme has the advantages of reducing noise and simplifying read-out connectivity (Fitzgerald et al., 2013). How did such biased representations develop? We next use modeling to infer the mechanisms by which a biased representation might be learned by LIP.

### Premises for a model of bias development in LIP

Category representation in LIP is a result of learning (Fitzgerald et al., 2013; Sarma et al., 2016). Before learning the task, the monkeys did not experience the stimuli as members of categories. Fig. 2A shows the rank ordering of LIP single neuron responses when passively viewing the same stimuli used in the match-to-category task, recorded from a monkey before it was trained on that task (Fitzgerald et al., 2013). Indeed, the neurons did not exhibit selectivity or representational bias for the “categories” which were not yet defined at that time.

**Figure 2.**
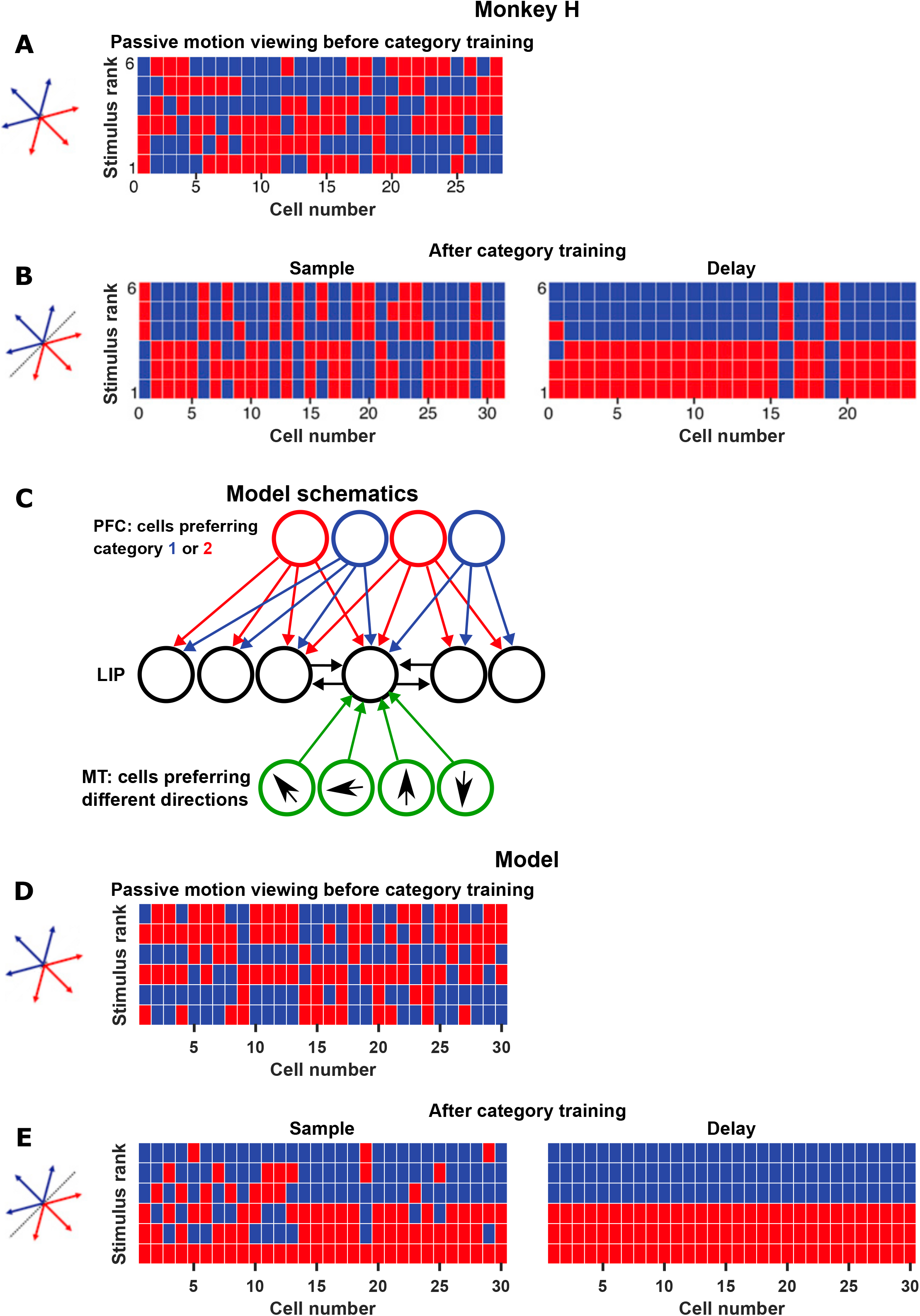
Model of LIP learning reproduces experimental data. (A) The selectivity of LIP cells from monkey H to the sample stimuli during passive viewing, before they were grouped into categories through training on the match-to-category task (Fig. 1A). Note that LIP did not exhibit category selectivity or bias before training. (B) The selectivity of LIP cells from monkey H after training. Replotted from the top panels of Fig. 1G. (C) Model schematics. In the model, we assumed that during training on the match-to-category task, PFC learned the categories first, and through plasticity at the PFC-to-LIP synapses, LIP cells acquired both category selectivity and biased representation. Equal numbers of PFC cells preferred categories 1 and 2, and they projected equally to LIP cells. LIP cells were recurrently connected with each other, and each received bottom-up inputs from MT neurons with a random set of preferred motion directions. See text for details. (D)-(E) Same as (A)-(B), but from a representative simulation of the model. The model reproduced the emergence of both category selectivity and biased representation. (A)-(B) adapted from Fitzgerald et al. (2013).

Which synaptic connections are responsible for the learning of categories in LIP? Unlike the classical visuospatial responses of LIP which do not require learning, category selectivity is a nonspatial response. Balan and Gottlieb (2009) inactivated LIP during three tasks in which LIP encoded three nonspatial variables: limb motor plans, estimated time, and reward expectation. LIP inactivation produced visuospatial deficits, but had no effect on the nonspatial aspects of performance related to the three variables. This suggests that LIP does not play a major functional role in processing nonspatial variables; instead, LIP may be simply reflecting the nonspatial signals in its external inputs, and using them for visuospatial processing.

Where do the external inputs carrying nonspatial signals come from? One likely candidate is PFC, an area reciprocally connected with LIP and known to encode abstract, cognitive factors and transmit them to posterior parietal cortex (Crowe et al., 2013). Recording during the same match-to-category task as Freedman and Assad (2006), Swaminathan and Freedman (2012) found that PFC also exhibited category selectivity.

Given these observations, we make the following assumptions, which serve as the premises of our model. As a monkey is being trained on the match-to-category task and learns to group stimuli into categories, PFC first acquires category selectivity, with approximately equal number of neurons preferring each category. As training progresses, the PFC-to-LIP synapses undergo Hebbian plasticity. Under conditions to be discussed below, such plasticity allows LIP to acquire both category selectivity and representational bias.

### Architecture and mechanisms of the model

We modeled a network of interconnected LIP neurons, which received bottom-up inputs from area MT and top-down inputs from PFC (Fig. 2C). The model PFC consisted of two populations with equal numbers of neurons, with each population preferring one of the two motion categories. Each LIP cell was innervated by equal numbers of PFC cells from each population. Starting from a naive state where LIP cells had no category selectivity (Fig. 2D), Hebbian plasticity at the PFC-to-LIP synapses resulted in the learning of category selectivity and representational bias (Fig. 2E). Below we present the details of the model and analyze its mechanisms, focusing first on the delay period.

During the delay period, no stimulus was shown and MT was silent, while PFC neurons had persistent delay activity. We modeled LIP activity during the delay with a standard linear firing rate equation (Dayan and Abbott, 2005):

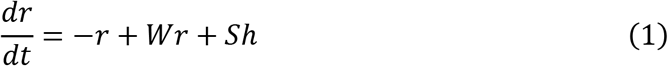

Here, *r* is a vector of the activity of LIP neurons, *W* is the LIP recurrent synaptic weight matrix, *S* is the PFC-to-LIP synaptic weight matrix, and *h* is the vector of PFC activity. LIP delay activity is then the steady-state response to persistent input from PFC:

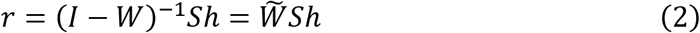

where *I* is the identity matrix, and we have defined 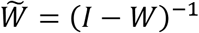.

In our model the only plastic synapses were the ones onto LIP neurons from PFC, that is, *S*. We initialized *S* with weak, random weights, and implemented lower and upper bounds on the weight of each synapse. We used a standard Hebbian plasticity rule to model the changes in *S*:

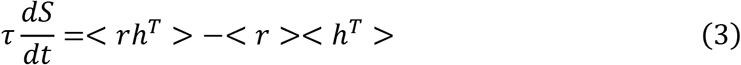

where *τ* is a time constant that controls the plasticity rate, and <•> denotes averages over all trials regardless of the stimulus presented. Thus, synapses potentiated (or depressed) when presynaptic firing in PFC and postsynaptic firing in LIP were correlated (or anti-correlated). Substituting (2) into (3), and assuming *S* changes slowly so that it can be regarded as constant when averaging over trials in (3), we obtain:

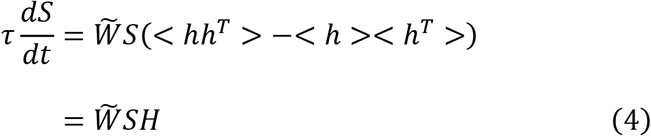

where we have defined the input covariance matrix *H* =< *hh*^*T*^ > −< *h* >< *h*^*T*^ >. If *r* is *N*-dimensional and *h* is *P*-dimensional, then 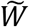 is *N* × *N, H* is *P* × *P*, and *S* is *N* × *P*.

(4) is a linear differential equation that can be understood as follows. The matrix 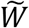can be characterized by its eigenvectors, which are the *N*-dimensional vectors 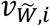 that it acts upon to return a multiple of the same vector: 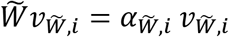, where 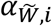 is the eigenvalue corresponding to the eigenvector 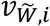 The subscript *i* specifies a particular eigenvector/eigenvalue pair, and for any generic square *N* × *N* matrix there are *N* such pairs, so that *i* takes values from 1 to *N*. The eigenvectors form a complete basis, meaning that any *N*-dimensional vector can be written as a weighted sum of these eigenvectors. Similarly *H* can be characterized by its *P* pairs of eigenvectors and eigenvalues, *v*_*H,j*_ and *α*_*H,j*_. Then the *N* × *P* matrices 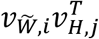 form a complete basis for *N* × *P* matrices, meaning that any *S* can be written as a weighted sum of such matrices: 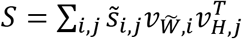 for a set of numbers 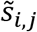. A given matrix 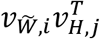 represents a synaptic connection pattern: the vector of synaptic weights from different PFC neurons onto any given LIP neuron is proportional to *v*_*H,j*_, with the proportionality constant for the *k*th LIP neuron being the *k*th element of 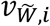

Substituting 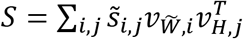 into equation (4) greatly simplifies the equation, because only the numbers 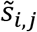 vary over time, while the eigenvectors are fixed; because the action of the matrices 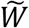 and *H* on a given submatrix 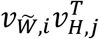 is just to multiply it by the corresponding eigenvalues; and because the resulting equation must be satisfied separately for each submatrix 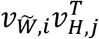 (Fig. 3A-B). As a result, equation (4) reduces to

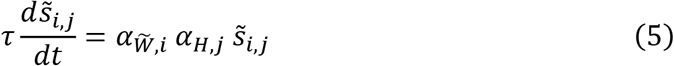

for all *i* and *j*. Starting with an initially random weight matrix *S*, the 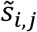 that grows the fastest will be that for which 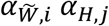 has the largest positive real part (Fig. 3C), and the structure of *S* will thus come to be dominated by the corresponding connection pattern 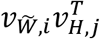.

**Figure 3.**
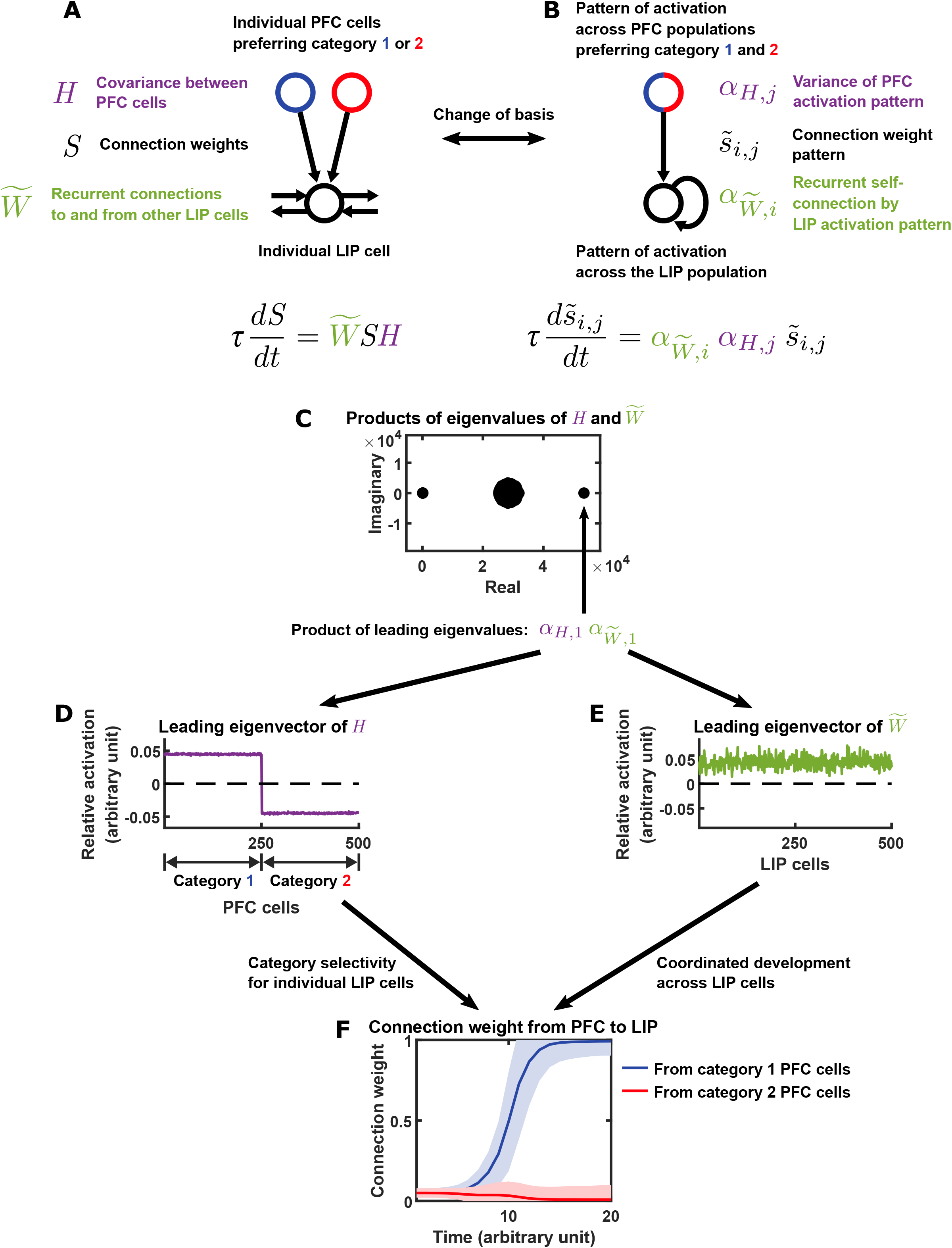
The mechanisms of category learning and bias development in the model. (A) Schematic of the model from the perspective of individual LIP and PFC cells. The weights of top-down PFC-to-LIP connections (*S*) develop based on the covariance of PFC cells (*H*) and the recurrent connections among LIP cells 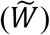. (B) A change of basis leads to a simplified view of the model, where single PFC activation patterns connect to single LIP activation patterns. Each PFC pattern is an eigenvector of *H* and the variance of the pattern is the eigenvalue *α*_*H,j*_ ; each LIP pattern is an eigenvector of 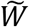 and the strength of its recurrent feedback is the eigenvalue 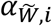. The connection 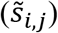 from a PFC pattern to an LIP pattern develops based on the variance of the PFC pattern (*α*_*H,j*_) and the feedback of the LIP pattern 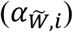, without interactions with other PFC and LIP patterns. See text for details. (C)-(F) Model mechanisms illustrated using an example simulation (the one from Figure 2D-E). (C) All pairwise products between the eigenvalues of *H* and the eigenvalues of 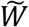. Each eigenvalue product is associated with a connection from a PFC pattern to an LIP pattern, whose weight changes at a rate proportional to the real part of the product. Therefore, the connection associated with the eigenvalue product with the largest positive real part would strengthen the fastest and dominate the connectivity pattern from PFC to LIP. This eigenvalue product is 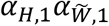, the product of the leading eigenvalues of *H* and 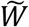. Their corresponding eigenvectors are illustrated in (D)-(E). (D) The leading eigenvector of *H*, representing the PFC activation pattern with the largest variance. PFC cells preferring the same category have the same sign (representing correlation between these cells), while PFC cells preferring different categories have opposite signs (representing anti-correlation between these cells). Strengthening the connection from this activation pattern to LIP leads to category selectivity: for a given LIP cell, all synapses from PFC cells preferring one category would strengthen, and all synapses from PFC cells preferring the other category would weaken. Note that this would not constrain different LIP cells to learn the same category preference. (E) The leading eigenvector of 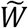, representing the LIP activation pattern most strongly amplified by the recurrent connectivity within LIP. All LIP cells have the same sign, representing coordinated activation caused by recurrent excitation between them. Such co-activation acts as positive feedback that causes different LIP cells to learn the same category preference, giving rise to biased representations. (F) Connection weights as a function of time, from PFC cells preferring category 1 (blue) and category 2 (red) onto LIP cells. Plotted are mean (± standard deviation) across all connections from PFC cells preferring a given category onto LIP cells. Note that in this simulation, connections from category 1 PFC cells strengthened while those from category 2 cells weakened, leading to a biased representation in LIP favoring category 1.

Thus, to understand the structure of changes in *S*, we need to characterize the “leading eigenvectors” of 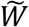 and *H*: the eigenvectors whose eigenvalues have the largest positive real part. We first examine *H*, followed by 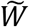. While the detailed structure of *H* depends on the specific activity statistics of the model PFC, one general feature of any reasonable PFC model is the anti-correlation between the two PFC populations preferring different categories. This is reflected in the structure of the leading eigenvector of *H* for the PFC in our model (Fig. 3D; see Supplementary Information for detailed analysis). This anti-correlation results in the dominant effect of *H* on the development of *S*: for a given LIP cell, all synapses from PFC cells preferring one category tend to increase in strength, and all synapses from PFC cells preferring the other category tend to decrease in strength. This gives rise to category selectivity for each LIP cell. Without the influence of the LIP recurrent connectivity 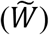, each LIP cell would learn its category preference randomly and independently, leading to an unbiased representation of the two categories. However, 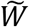 can act to coordinate the learning of category preference among LIP cells. If the LIP recurrent connectivity has stronger excitation than inhibition, then the leading eigenvector of 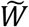 consists of elements with the same sign (Fig. 3E). This means that LIP neurons tend to activate together, leading to a positive feedback loop where they learn the same category preference, resulting in a biased representation. The stronger the recurrent excitation (and the weaker the recurrent inhibition), the more likely different LIP neurons are to learn the same category preference (see Discussion for a discussion on the variability of bias across experimental animals and across simulations).

Intuitively, the effects of PFC input covariance and LIP recurrent connectivity on synaptic development can be summarized as follows. Consider a single LIP cell before training, which receives weak and roughly equal inputs from PFC populations 1 and 2 that each prefers one of categories. During early training, the random synaptic connections from the two PFC populations and their noisy firing could lead one PFC population, say population 1, to activate the LIP cell slightly more than the other PFC population. This would lead to slight potentiation of the population 1 synapses onto the LIP cell, and a slight depression of the population 2 synapses onto it. Because of these synaptic changes, later on in training, PFC population 1 would activate the LIP cell more, leading to more potentiation of their synapses. Conversely, the synapses from PFC population 2 would depress. This cycle of positive feedback eventually would lead to large weights at population 1 synapses and small weights at population 2 synapses onto this LIP cell, making it prefer category 1. Now consider other LIP cells in the network. When the one LIP cell first starts to weakly prefer category 1, it excites other inter-connected LIP cells, biasing them to also fire more in response to category 1. This leads to potentiated synaptic weights from PFC population 1 to all these LIP cells, which recurrently excite each other to further increase their response to category 1. This second cycle of positive feedback across LIP cells eventually leads to most of them preferring the same category (Fig. 3F).

In our model, the bottom-up sensory projections from MT to LIP were taken to be hard-wired and not plastic during category training. In particular, each LIP cell received input from MT cells preferring motion directions that were random with respect to the arbitrary categories. Thus, in the model, visual input from MT to LIP during the sample period, which were absent during the delay period, weakened the bias in LIP compared to the delay period (Fig. 2E).

### Development of bias after re-definition of categories

Our model can also reproduce the re-development of bias after re-definition of categories, which was observed by Fitzgerald et al. (2013). Freedman and Assad (2006) had trained monkeys on the match-to-category task, with 12 directions belonging to 2 categories. As the monkeys learned the categories, biased representations developed in LIP (Fig. 4A and C). Then, the categories were re-defined, such that the new category boundaries were orthogonal to the old ones, dividing the 12 directions into 2 new categories (Fig. 4A-D). Since the old and new category boundaries were orthogonal, the old category representation in LIP would translate to neither category selectivity nor category bias under the new definitions. However, after the monkeys learned the new categories, the neural representation in LIP changed, such that neurons showed both category selectivity and biased representation for the new categories (Fig. 4B and D).

**Figure 4.**
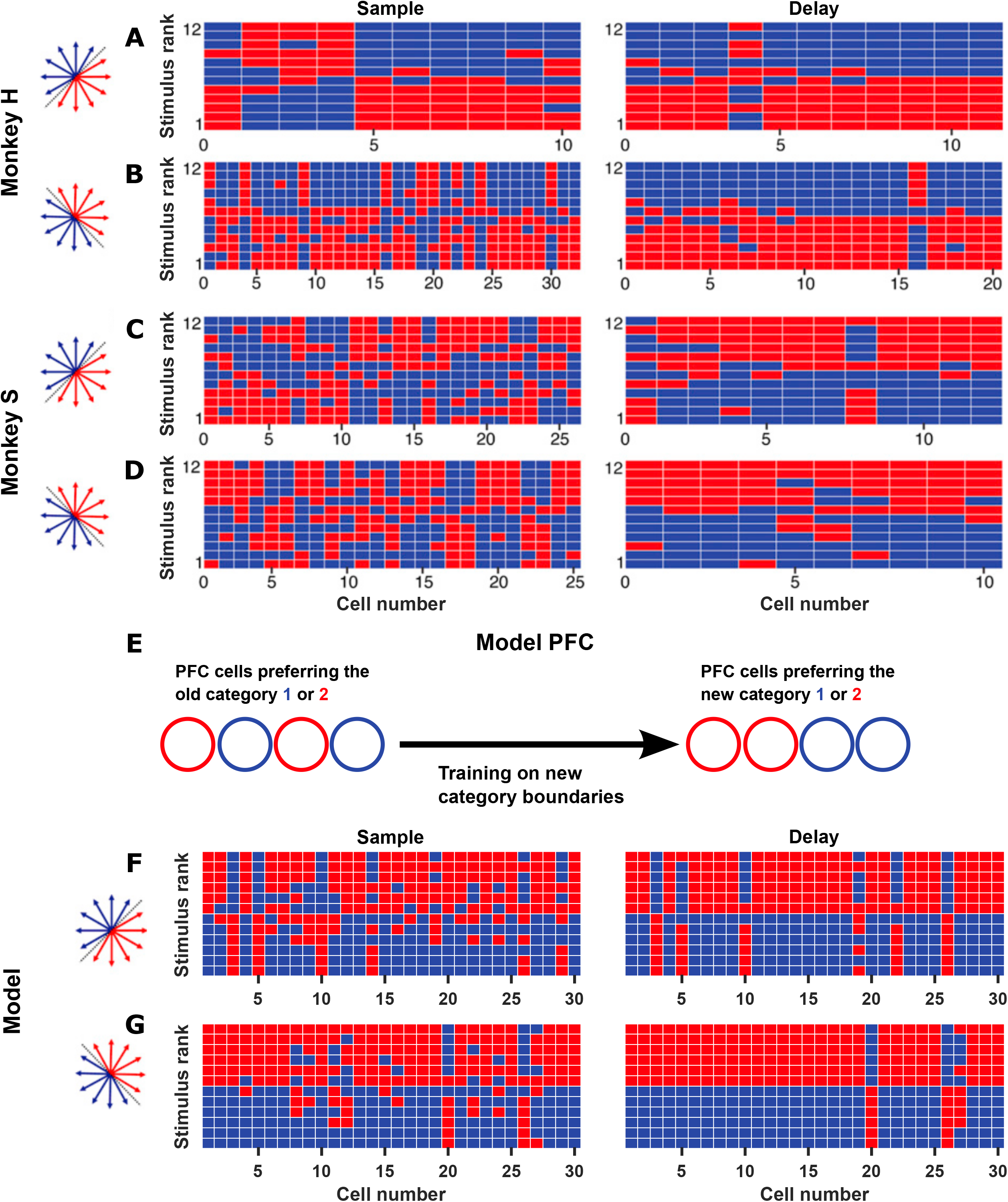
Model reproduces re-emergence of biased representation after re-definition of categories. (A)-(D) Monkeys H and S were first trained to group 12 motion directions into two categories, and LIP learned biased representations of the categories (A) and (C). The monkeys were then trained to group the same motion stimuli into two new categories, and LIP again learned biased representations of the redefined categories (B) and (D). (E) The model PFC had an equal number of neurons encoding each of the original categories. After training on the original categories, the categories were re-defined: half of each PFC population was assigned to each new category. Training then proceeded to modify the PFC-to-LIP connection weights given the new category representation in PFC. (F)-(G) The model reproduced biased representations of a first set of categories (F), as well as re-development of bias after re-definition of the categories (G). (A)-(D) adapted from Fitzgerald et al. (2013).

To model this phenomenon, we made further assumptions on the neural representation in PFC as animals learned new category definitions (Fig. 4E). As the old and new category boundaries were orthogonal, for the PFC cells that encoded a given old category, a random half was assigned to each of the new categories, without altering the PFC-to-LIP connection weights learned during the first category training. Training then proceeded to modify the PFC-to-LIP connection weights given the new representation in PFC.

Training on the first set of categories resulted in category selectivity and bias in this model, as expected (Fig. 4F). During the re-training on new categories, the change in the PFC representation changed the covariance matrix *H*, which drove LIP neurons to again develop category selectivity and bias for the new categories, by the same mechanism as explained in the previous section (Fig. 4G).

### Development of bias for three categories of shapes

In another set of match-to-category experiments, Fitzgerald et al. (2011) trained monkeys to categorize six abstract shapes into three categories (Fig. 5A-B). In this experiment, a category bias developed in one of the monkeys (Fig. 5A; see Discussion for a discussion on the variability of bias across experimental animals).

**Figure 5.**
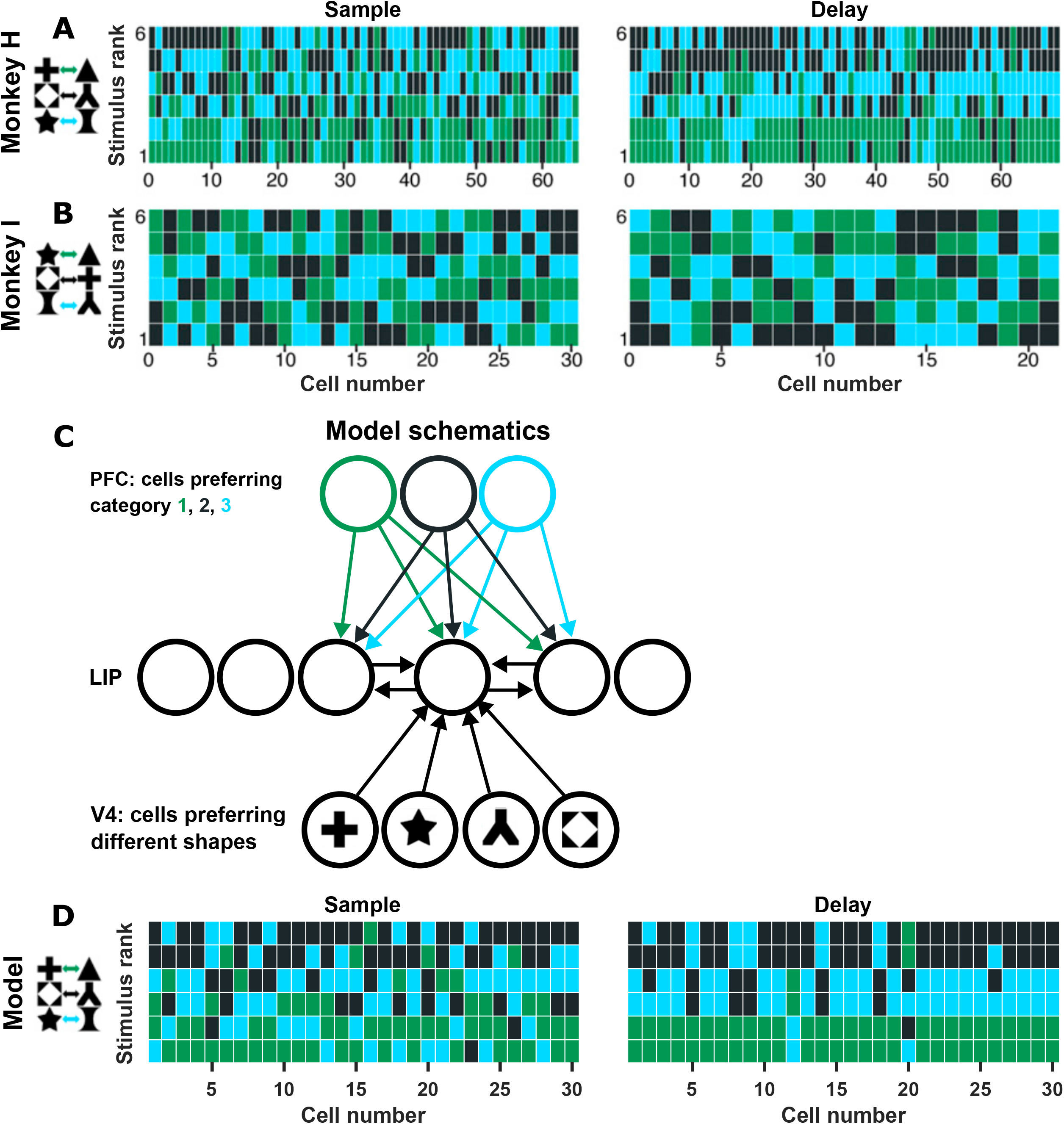
Model reproduces the biased representation of three categories of abstract shapes. (A)-(B) Monkeys H (A) and I (B) were trained on a match-to-category task with identical structure as in Fig. 1A but different stimuli and categories: six abstract shapes grouped into three categories. Monkey H (A) learned a biased representation in this task. (C) Model schematics. Same as the model in Fig. 2C, except that there were three equal PFC populations encoding the three categories, and that the bottom-up inputs to LIP in this task came from V4 shape-selective neurons. (D) Model reproduced biased representations of three shape categories. (A)-(B) adapted from Fitzgerald et al. (2013).

To model LIP learning in this task, we assumed that the sensory inputs encoding the shapes came to LIP from V4, with each LIP cell receiving inputs from V4 cells preferring a random set of shapes. Furthermore, we assumed that after the animals learned the categories, PFC contained three equal populations of cells preferring each of the three categories, which sent equal projections to LIP (Fig. 5C).

In this case, the PFC covariance structure was similar to the two-category case analyzed above, where PFC cells preferring the same category were correlated, while PFC cells preferring different categories were anti-correlated. The effect of this covariance structure on PFC-to-LIP synaptic plasticity led to LIP cells developing differential responses to the three categories, making them category-selective, as in the direction category tasks above (see Supplementary Information for a discussion on the effects of connection weight constraints on category selectivity). The recurrent connectivity in LIP again led to LIP cells learning the same category preference, resulting in biased representations (Fig. 5D).

### Spatial clustering of LIP category preferences

In our model, LIP cells gave positive feedback to each other during learning via their excitatory recurrent connections, making connected cells learn the same category preference. In the brain, physically nearby cells are more heavily recurrently connected, so our model suggests that there might be spatial clustering of category preferences. To examine this, we modeled an LIP network on a 2D cortical surface. The LIP cortical surface is known to contain rough topological maps of visual space: neurons whose RFs are nearby are more likely to be located close to each other on the cortical surface (Blatt et al., 1990; Patel et al., 2010). In our simple model, there was an exact topological correspondence between neurons’ location on the cortical surface and their RF positions. Thus, the connection probability between LIP neurons was a decreasing function of the distance between their RF positions. In this model, category selectivity developed as in the main model above. Furthermore, we modeled long-range connections between excitatory cells to inhibitory cells within LIP. This encouraged cells located away from each other to learn different category preferences (as opposed to a strongly biased representation where most cells prefer the same category). In this model, we observed spatial clustering of category preferences (Fig. 6A).

**Figure 6.**
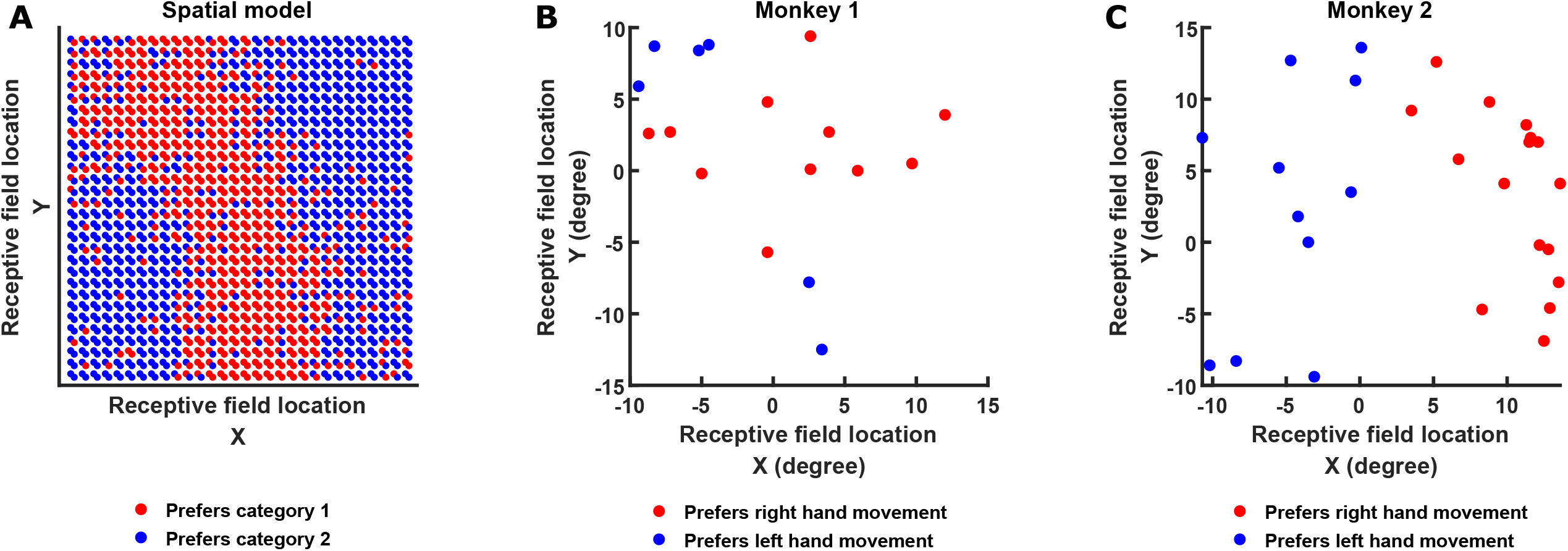
Data confirms model prediction that cells with the same preference should have spatially clustered receptive fields. (A) We modeled a LIP network on a 2D cortical surface, where neurons had a topological representation of visual space. Connection probability between LIP neurons fell off with distance on the surface, and thus with distance between their RFs in visual space. Shown are the category preference of cells in a model network after category training, plotted as a function of their RF location, showing that patches of cells with the same category preference developed. (B)-(C) In a visual discrimination task where decisions were reported using either the left or right hand, Oristaglio et al. (2006) found that LIP cells had hand selectivity independent of visuospatial information. Plotting the hand preference of LIP cells as a function of their RF location confirms our model prediction that cells with the same preference would have RFs that cluster in space. Data from two monkeys are shown in (B) and (C).

We wished to test our model prediction that cells with the same category preferences would have spatially clustering RFs. The Fitzgerald et al. (2013) datasets are not a good test for this prediction, since the strong biases in those datasets means that almost all cells had the same category preference. Instead, we turned to an experiment by Oristaglio et al. (2006), where LIP exhibited weaker biases of response preferences. Briefly, their behavioral task is as follows. A monkey fixated a central fixation spot and held two bars with its two hands. There were four figure-8s on the screen, and after ∼500 ms of fixation, two line segments were removed from each figure-8. This resulted in one of the four stimuli turning into either a leftward-facing or rightward-facing letter “E”, which was the task-relevant cue for the monkey, while the other stimuli turned into task-irrelevant distractors. The monkey was rewarded if it released its right hand when the “E” was rightward-facing and released its left hand when the “E” was leftward-facing, while maintaining fixation. In this task, LIP neurons encoded the hand that was being used to release a bar. This hand selectivity was dissociated from selectivity for the identity of the stimulus or the position of the hand (Oristaglio et al., 2006).

This hand selectivity was similar to the category selectivity observed by Fitzgerald et al. (2013) in that they were both “non-spatial” response properties, as opposed to the classic spatial responses of LIP neurons, including visual response, delay response, and saccadic responses. Furthermore, hand selectivity was likely learned through training, like category selectivity. Thus, we hypothesized that hand selectivity developed in LIP through the same mechanisms we proposed for category selectivity. Thus, we predicted hand preference of LIP cells would show spatial clustering. Indeed, this was what we observed: cells preferring left hand movements tended to have RFs located near other left-hand-preferring cells, and similarly for right-hand-preferring cells; the RFs of cells with different hand preferences were separated into spatial clusters (Fig. 6B-C).

## Discussion

### Further predictions and experimental tests

In addition to the prediction of RF spatial clustering by preferred category, which we verified in the data of Oristaglio et al. (2006), the model offers several additional predictions. One premise of our model is that category learning in LIP is guided by PFC. In our model, this scenario was simplified, such that category learning in PFC was completed before category learning started in LIP. In reality, category learning in the two areas likely proceeds in parallel, but with LIP lagging and being guided by PFC. This can be tested by simultaneous recording from LIP and PFC during the process of training on the match-to-category task.

In the data of Fitzgerald et al. (2013), category bias was weaker or absent during the sample period (Fig. 1G, Fig. 4A-D, Fig. 5A-B). In our model, connections from MT or V4 to LIP were random and not plastic; thus, these random bottom-up input weakened the bias in LIP during the sample period compared to the delay period (Fig. 2E, Fig. 4F-G, Fig. 5D). This leads to the prediction that, if PFC were inactivated during the sample period, any bias in LIP would disappear, because LIP would be mainly driven by random bottom-up inputs. A simpler prediction to test is that, if PFC is found not to be category selective during passive viewing outside of the task, then LIP stimulus preferences during such passive viewing should be uncorrelated with their preferences during the delay period of task performance.

Fitzgerald et al. (2013) suggested that biased representation might be a neural strategy for encoding discrete variables, such as categories, and that LIP neurons might encode continuous variables with more unbiased representations. However, the learning mechanisms we proposed is compatible with LIP learning biased representations of continuous variables. In tasks where animals compare the magnitude of continuous stimuli, neurons can encode the continuous stimulus magnitude. For example, Ferrera et al. (2009) trained animals to compare the speed of moving dots, and found two roughly equal populations of FEF neurons with monotonic tuning functions for speed, with one population preferring faster speeds and another preferring slower speeds. If such prefrontal populations could guide the learning of continuous variables in LIP as we proposed above for the learning of categories, we predict that LIP would learn biased representations for the continuous variable. For example, Fig. S1 shows a simulation where almost all LIP neurons learned to prefer larger magnitudes of the continuous variable. In Fig. S1A we propose an experimental task where this prediction might be tested.

### Difference in degree of bias across animals

Among the animals analyzed in Fitzgerald et al. (2013), not every animal developed a biased representation (Monkey I did not develop a bias in the shape-category task and had a weaker bias in the direction category task; Fig. 1G and 5B). In the model, there was also stochasticity in the learning outcome, ranging from no bias to strong bias (results not shown). The learning outcome depends on the random initialization of PFC-to-LIP weights, as well as on how strongly the LIP recurrent connectivity amplifies its most strongly amplified activity pattern (the leading eigenvector of 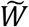) relative to other activity patterns. Tuning model parameters to more strongly amplify the leading eigenvector (e.g., by increasing recurrent excitation relative to inhibition) makes a biased representation more likely to develop. Given the limitations on sample sizes in monkey experiments, it is currently unknown whether there is substantial variability in the degree of bias across animals. Furthermore, it is not known whether any differences in the degree of bias is causally related to differences in LIP connectivity (e.g., the strength of recurrent excitation). Whether or not bias develops, and to whatever degree it develops, we predict that cells of the same category preference should have spatially clustered RFs.

### Why does LIP have connectivity that favors biased representations?

Our model suggests that biased representations develop in LIP due to the structure of its recurrent connectivity. We believe the ability of LIP to learn biased representations is indicative of its wider role in visuospatial cognition (Gottlieb, 2007; Gottlieb et al., 2009; Gottlieb and Snyder, 2010). Specifically, this role of LIP is to learn to encode the significance of visual stimuli, in order to guide attentional allocation and eye movements. We elaborate on this idea below.

The biased representation of abstract categories examined here are not intuitive— it is not obvious why LIP neurons would all prefer one non-spatial stimulus feature over others. However, other “biases” that LIP neurons exhibit are intuitive and taken for granted. For example, in tasks where different visual stimuli are associated with different amounts or probabilities of reward, almost all LIP neurons prefer stimuli predicting larger rewards (Coe et al., 2002; Dorris and Glimcher, 2004; Sugrue et al., 2004; Peck et al., 2009). Similarly, almost all LIP neurons prefer task-relevant stimuli over irrelevant ones (Oristaglio et al., 2006). We suggest that such “biases” for reward-predicting stimuli could have developed in LIP through the same mechanisms that we proposed for the development of biased category representations. As long as reward learning in LIP is guided by areas where at least a small majority of cells prefer larger rewards (e.g., FEF or dlPFC, discussed more below), LIP could reliably learn a strongly biased representation that prefers larger rewards. Thus, when faced with reward-predicting stimuli, these learning mechanisms allow LIP to identify behaviorally significant stimuli, allowing attention to be directed towards them covertly or overtly.

Interestingly, in FEF and dlPFC, areas that we hypothesize guide LIP in learning the significance of stimuli, only a small majority of neurons prefers stimuli associated with higher rewards (Leon and Shadlen, 1999; Coe et al., 2002; Roesch and Olsen, 2003; Pan et al., 2008; Kennerly and Wallis, 2009; Teichert et al., 2014). That is, PFC contains neural populations preferring stimuli predicting larger rewards as well as preferring smaller rewards. It is possible that this is because PFC is involved with encoding all stimuli regardless of their current behavioral significance, allowing flexible behavior in environments where the significance of stimuli can change. Through plasticity at the PFC-to-LIP synapses, LIP is able to learn the *current* significance of stimuli from PFC and direct attentional processes accordingly. Thus, through learning, LIP may become heavily involved in mediating the currently relevant behavioral task, while PFC remains more involved in general cognitive computations and less involved in any specific task (Swaminathan and Freedman, 2012).

In summary, our model identifies a possible mechanism by which the recurrent connectivity structure of LIP allows it to learn abstract representations, which enables it to assign behavioral relevance to stimuli in order to guide behavior.

## Methods

### LIP connectivity

To model the results of Freedman and Assad (2006) and Fitzgerald et al. (2011), we modeled a LIP network with *N* cells, half of them excitatory (E) and half of them inhibitory (I), with matrix *W* (*N* × *N*) describing their recurrent connectivity. Recurrent connections in LIP are random and sparse with connection probability *p* (a random proportion *p* of all possible excitatory connections were given nonzero weights, and the same was done for inhibitory connections; autapses were not allowed). The nonzero weights were drawn from Gaussian distributions whose parameters depended on the cell types of the pre- and post-synaptic cells: the mean weights for E-to-E, E-to-I, I-to-E, and I-to-I connections were *w*_*EE*_, *w*_*IE*_, *w*_*EI*_, and *w*_*II*_, respectively, the standard deviations were the absolute values of the respective means scaled by a factor 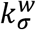. Randomly drawn negative weights excitatory connections were set to zero, and similarly positive weights for inhibitory weights were set to zero; 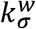 was set such that the probability for such events was very low.

For our model of an LIP network on a 2D surface, we modeled a *N*_*s*_ × *N*_*s*_ grid with a pair of E and I cells at each grid position. The connection probability *p*_*ij*_ between two different cells *i* and *j* falls off as a function of the distance *d* between them: 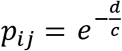, where the connection range parameter *c* takes the value *c*_*EE*_, *c*_*IE*_, *c*_*EI*_, or *c*_*II*_ for E-to-E, E-to-I, I-to-E, or I-to-I connections, respectively. Connection weights were drawn from Gaussian distributions as above.

### Top-down connectivity

We modeled top-down inputs to LIP from *N*_*f*_ E cells in PFC, consisting of equal numbers of cells preferring each category (three categories for the shape category experiment, two categories in all other cases; in the case of the continuous stimulus model, equal numbers of cells preferred larger and smaller stimulus values). Top-down connections from PFC were represented by the matrix *S* (*N* × *N*_*f*_). Connections were sparse with connection probability *p* (each LIP cell received projections from a random proportion *p* of the PFC cells preferring each category). For the connected cells, the initial projection weights were independently drawn from a uniform distribution on (0, 0.1). These top-down projections were plastic and were responsible for learning, as described below.

### PFC responses and covariance

For each stimulus category, we generated 1000 instances of PFC activity patterns *h* (*N*_*f*_ × 1): in each instance, each PFC cell’s activity was drawn from a Gaussian distribution whose mean was 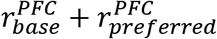 for the preferred category or 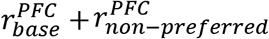 for non-preferred categories, and whose standard deviation was the mean scaled by 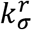. Randomly drawn negative activity were set to zero; 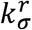 was set such that the probability for such events was very low. The PFC covariance matrix *H* was then estimated from these instances of activity patterns.

We also generated PFC spontaneous activity for the purpose of evaluating LIP responses before training (described below), as follows. Each PFC cell’s activity was drawn from a Gaussian distribution whose mean was 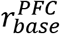 and whose standard deviation was the mean scaled by 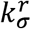 (randomly drawn negative activity were set to zero).

For the version of the model with a continuous stimulus (Fig. S1), each PFC cell was randomly assigned a response range 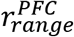 from a Gaussian distribution with mean *μ*_*range*_ and standard deviation *σ*_*range*_. Then we generated 1000 stimuli values from a uniform distribution from 0 to 1. For a stimulus value *α*, the response of a given PFC cell was 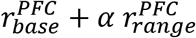 if the cell preferred larger stimulus values, or 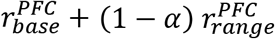 if it preferred smaller stimulus values. The PFC covariance matrix *H* was then estimated from the responses to these 1000 stimuli.

### Synaptic plasticity

We modeled the changes in the PFC-to-LIP connectivity matrix *S* with a standard Hebbian plasticity rule [equation (4) in Results]. Equation (4) was simulated using Euler’s method from *t* = 0 to *t* = 30. During learning, the weight between each connected PFC-LIP cell pair was bounded between 0 and 1, and the weight between each unconnected PFC-LIP cell pair was fixed at 0.

For the category re-definition experiment (Fig. 4), after the original category definition was learned, the categories were redefined for PFC cells without modifying *S*: half of each PFC population was assigned to each new category—e.g., half of the PFC population for the old category 1 now preferred the new category 1, and half now preferred the new category 2. Learning then proceeded as described above.

### Evaluating LIP responses

We evaluated the responses of model LIP networks to top-down and bottom-up inputs, before and after training. The steady-state LIP response was calculated as (*I* − *W*)^−1^*S*_*top*+*bottom*_ *h*_*top*+*bottom*_, where *S*_*top*+*bottom*_ = (*S S*_*bottom*−*up*_) consisted of the top-down projection matrix *S* and the bottom-up projection matrix *S*_*bottom*−*up*_, and 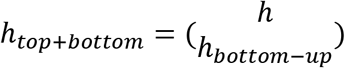 consisted of the PFC input *h* and the bottom-up input *h*_*bottom*−*up*_. *S* and *h* have been described above, and *S*_*bottom*−*up*_ and *h*_*bottom*−*up*_ are described below.

We modeled bottom-up visual sensory inputs to LIP from MT for the direction category experiments or from V4 for the shape category experiment. We modeled a total of *N*_*b*_ E cells from a given visual sensory area, consisting of equal-sized populations preferring each of the stimuli used in a given experiment. To construct *S*_*bottom*−*up*_ (*N* × *N*_*b*_), for each LIP cell, we randomly selected from a number *n*_*p*_ of the sensory populations to project to that cell (for each LIP cell, *n*_*p*_ is drawn from a discrete uniform distribution from *n*_*lower*_ to *n*_*upper*_). For each selected sensory population, a random proportion *p* of its cells were chosen to project to the LIP cell. These bottom-up sensory projections were taken to be hard-wired and not plastic, and they all had a weight of 1.

*h*_*bottom*−*up*_ (*N*_*b*_ × 1), the bottom-up inputs, was modeled as follows depending on whether it represented inputs from MT or V4. When modeling inputs during the delay period (when there was no visual stimulus), the mean spontaneous activity of each sensory cell was 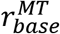or 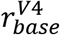 depending on whether it was modeling MT or V4. When modeling inputs during visual stimulus presentation, for the direction category experiments, the mean activity of an MT cell with preferred direction *θ*_*preferred*_ in response to a stimulus with direction *θ* was 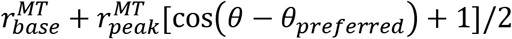. For the shape category experiment, the mean activity of a V4 cell was 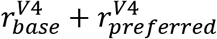 in response to its preferred shape and 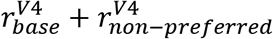 to its non-preferred shapes. To generate an instance of *h*_*bottom*−*up*_ for a given stimulus or for the delay period, the activity of each cell was drawn from a Gaussian distribution with the above means and with standard deviations that were the respective means scaled by 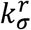 (randomly drawn negative activity were set to zero).

To examine the responses of the model LIP network from a given simulation (Figs. 2, 4, 5), we generated an instance of *h* in response to each category, an instance of *h* for PFC spontaneous activity, an instance of *h*_*bottom*−*up*_ in response to each stimulus, and an instance of *h*_*bottom*−*up*_ for MT or V4 spontaneous activity. For evaluating LIP response during the sample period, *h*_*top*+*bottom*_ was formed by combining PFC category response with the appropriate bottom-up visual response. For evaluating LIP response during the delay period, *h*_*top*+*bottom*_ was formed by combining PFC category response with the MT/V4 spontaneous activity. For evaluating LIP response to the direction stimuli before training, *h*_*top*+*bottom*_ was formed by combining PFC spontaneous activity with MT visual response. In all cases, the responses of all cells in the LIP network were calculated as (*I* − *W*)^−1^*S*_*top*+*bottom*_ *h*_*top*+*bottom*_, and responses of a random subset of cells were ranked and visualized. When evaluating LIP response for the category re-definition experiment (Fig. 4), different instances of PFC category responses were used for the first and second category definitions, while the same instances of MT visual and spontaneous activity were used.

The tuning curves in Fig. S1B-C were calculated as follows. As described above, for a stimulus value *α*, the response of a given PFC cell was 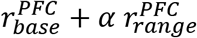 if the cell preferred larger stimulus values, or 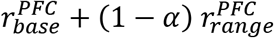 if it preferred smaller stimulus values. Using *h*_*α*=0_ and *h*_*α*=1_ to denote PFC responses to *α* = 0 and *α* = 1, we calculated the corresponding LIP response as (*I* − *W*)^−1^*Sh*_*α*=0_ and (*I* − *W*)^−1^*Sh*_*α*=1_.

(*I* − *W*)^−1^*Sh*_*α*=0_ and (*I* − *W*)^−1^*Sh*_*α*=1_ were then z-scored together, and LIP responses to *α* between 0 and 1 were linearly interpolated.

### Parameters

Model of motion category experiment:

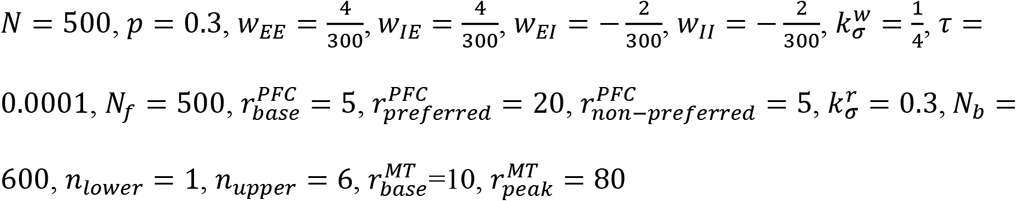

Model of motion category re-definition experiment:

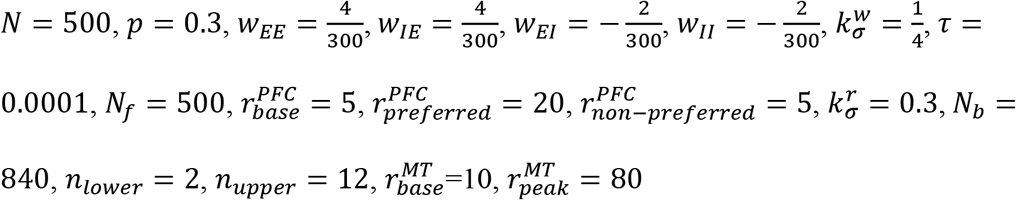

Model of shape category experiment:

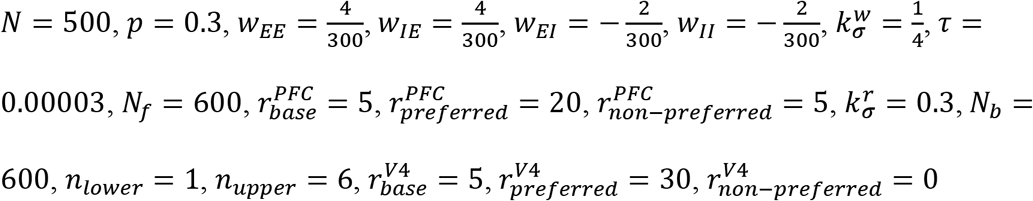

Model of LIP network on 2D surface:

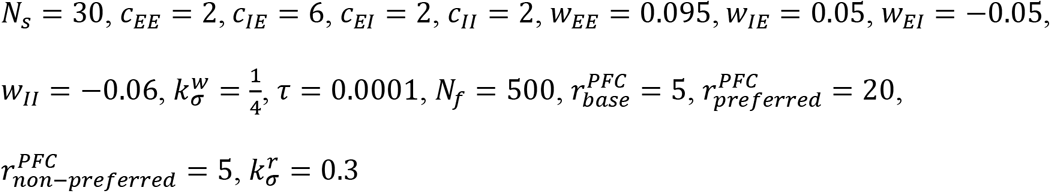

Model with continuous stimulus:

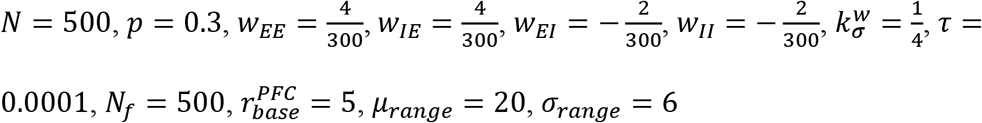

### Analysis of the Oristaglio et al. (2006) data

For each cell, spike counts from -200 to 0 ms before bar release on left hand release trials were compared with those on right hand release trials, using a two-sample t test. A cell with a *p* value of less than 0.05 was considered hand-encoding.

## Supplementary Information

### Analysis of the covariance structure of the model PFC

Here we analyze the covariance structure of the model PFC, *H*, in more detail. Using h^*i*^ to denote PFC population activity on trials where category *i* was presented,

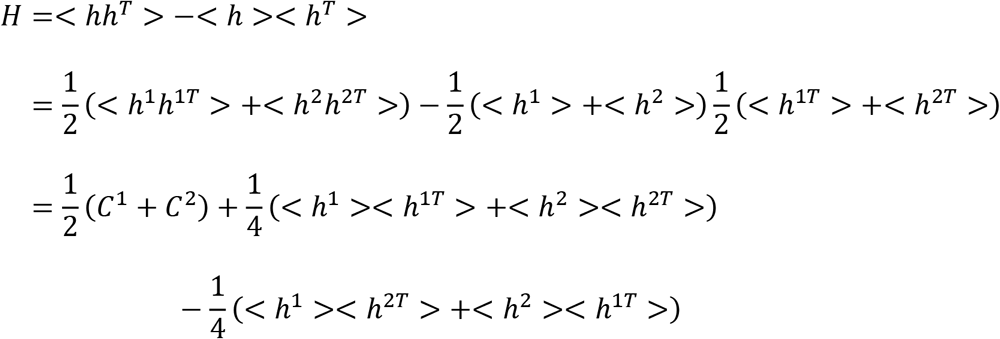

Here *C*^*i*^ denotes the PFC covariance matrix on trials where category *i* was presented. In our model, the responses of different PFC cells to the same category are independent, so we can simplify *H*:

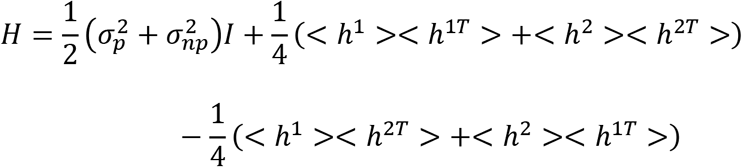

where 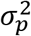 and 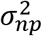 denote the variance across PFC cells of response to the preferred and non-preferred categories, respectively, and *I* is the identity matrix. We take the elements of the *h* vectors to be ordered such that the first 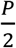 elements correspond to the PFC cells that prefer one category and the next 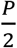 elements to the cells that prefer the other category. Using *m*_*p*_ and *m*_*np*_ to denote the mean response of a PFC cell to its preferred and non-preferred categories, respectively, and *A* to denote the square matrix of all ones with half the number of rows and columns as *H*, we can rewrite *H* as

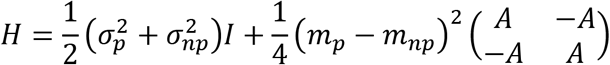

The first term represents the variances of individual PFC cells, and the second term represents the anti-correlation between the two PFC populations.

### Effects of connection weight constraints on learning in the three-category model

In typical simulations of the three-category model, connection weights from two of the PFC populations would potentiate together or depress together, with different rates for the two populations. At the same time, connection weights from the third population would change in the opposite direction compared to the first two populations. This results in three levels of LIP activity in response to the three categories. However, because our model implemented hard upper and lower bounds on connection weights, if simulations were allowed to proceed for long enough, connection weight from the first two populations would saturate at the same level, meaning that LIP would no longer have different levels of activity in response to these two categories. In our three-category model, plasticity rate was set such that the connection weights did not reach saturation by the end of learning, so that LIP developed three distinct levels of activity for the three categories. Alternative choices of connection weight constraints could allow LIP to retain three distinct levels of activity even at saturation: e.g., constraints where each weight decays at a rate proportional to its current value (Miller and MacKay, 1994). It’s worth noting that experimental data from the three-category experiment did show similar activity levels in LIP for two of the three categories (see Fig. 2H in Fitzgerald et al., 2013), and it’s possible that weight saturation played a role.

## Supplemental Information

**Figure S1.**
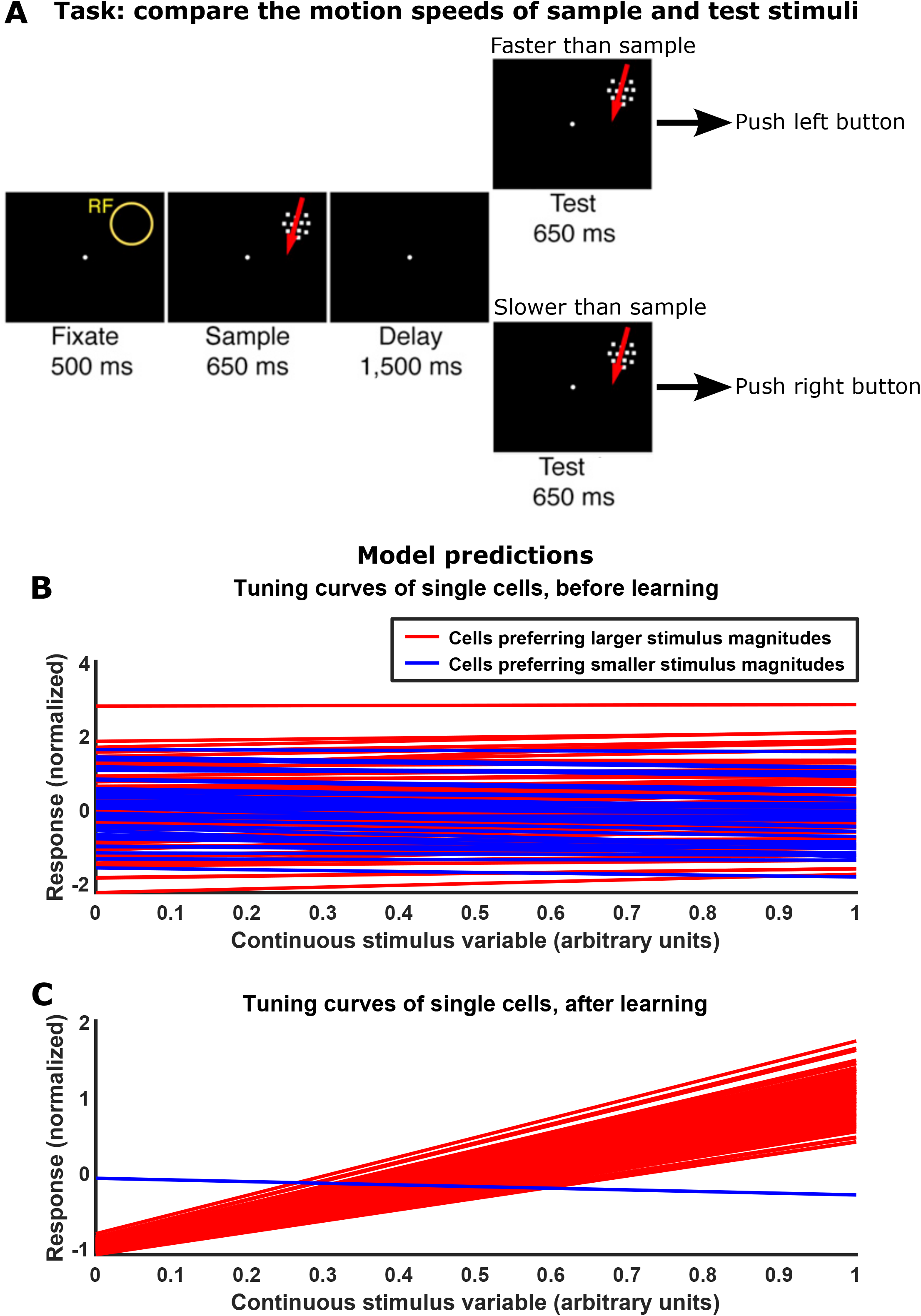
Model predicts biased representation of continuous stimulus variables. (A) A proposed speed comparison task modified from the match-to-category task in Fig. 1A. All sample and test stimuli are coherent dot motion in the same direction; the monkey is required to compare the speed of the sample and test stimuli. Figure adapted from Fitzgerald et al. (2013). (B)-(C) Model predicts that LIP would learn biased representations of continuous stimuli, such as motion speed in the task in (A), where almost all cells would prefer larger magnitudes or almost all cells would prefer smaller magnitudes of the stimulus. Plotted are the tuning curves of model LIP cells before (B) and after (C) learning, with each line being the tuning curve of one cell, colored according to whether the cell preferred smaller or larger stimulus values.

